# Gene networks in cancer are biased by aneuploidies and sample impurities

**DOI:** 10.1101/752816

**Authors:** Michael Schubert, Maria Colomé-Tatché, Floris Foijer

## Abstract

Gene regulatory network inference is a standard technique for obtaining structured regulatory information from, among other data sources, gene expression measurements. Methods performing this task have been extensively evaluated on synthetic, and to a lesser extent real data sets. They are often applied to gene expression of human cancers. However, in contrast to the evaluations, these data sets often contain fewer samples, more potential regulatory links, and are biased by copy number aberrations as well as cell mixtures and sample impurities. Here, we take networks inferred from TCGA cohorts as an example to show that (1) transcription factor annotations are essential to obtaining reliable networks, and (2) even when taking these into account, we should expect between 20 and 80% of edges to be caused by copy number changes and cell mixtures rather than transcription factor regulation.

## 1. Introduction

Gene Regulatory Network (GRN) inference describes the process of identifying regulator-target relationships from experimental molecular data. These data can be protein-protein interactions (often referred to as interaction networks) or protein-DNA binding (obtained by chromatin immunoprecipitation and sequencing, or ChIP-seq), but is most commonly gene expression data (obtained by microarrays or more recently RNA-seq). For gene expression data, the rationale behind the approach is that if a transcription factor (TF) is more highly expressed, it is also likely to be more active and mediate a higher downstream expression of its target genes (TGs). While this ignores potential post-translational modifications that may also influence a transcription factor’s activity, as well as epigenetic marks at the enhancer and promoter sites of target genes, GRN inference methods have been shown to be useful in elucidating transcriptional programmes in a variety of contexts ^1–7^.

There are different kinds of data that we can use to infer networks from. For instance, we can follow a perturbation over time (time-course networks), or take multiple snapshots of the same underlying system in different states. The latter is referred to as observational (meaning comparing different samples) steady-state networks ^8^, which often occur when we for instance measure gene expression in a yeast strain with different growth conditions or cancer patients across a cohort of the same tumor type.

Each of these use cases require different assumptions and hence tools that are specialized for this kind of data. Here, we focus on steady-state observational networks. In this case, we assume the underlying regulatory structure to be the same or at least its differences small enough so we can ignore them. This is likely true when e.g. a mutation in a signaling molecule activates a certain part of a downstream GRN, but it will not be if a transcription factor loses its affinity to its target genes or a subset thereof. While previous studies have shown this to happen for some genes (reviewed in ^9^), observational GRN inference methods assume that this will not change the overall correlation structure across many samples.

Methods that have been developed for observational GRNs can roughly be classified by the theoretical framework they use in order to infer regulatory relationships. The classical approaches come from information theory and employ some kind of mutual information, or correlation and regression-based approaches (classification and theoretical background have been reviewed before ^8^). These tools have been continuously developed, but more recently the focus has also shifted more to machine learning methods such as random forest and neural networks (recent overview of methods reviewed in ^10^).

These network inference methods have been extensively evaluated e.g. in the Dialogue of Reverse Engineering and Assessment of Methods (DREAM) competitions ^11^ and many more comparisons on smaller scale ^12–18^. They have provided many biological insights, and have been particularly useful to elucidate mechanisms of pathogenicity in human diseases such as cancer ^1,2,19–24^. However, there is a disconnect between evaluation in often relatively simple systems (synthetic networks or GRNs in *E. coli* and yeast) and their application to much more complex mammalian systems.

One application where this disconnect is particularly striking is human cancer, because (a) individual patients harbor different chromosomal aberrations ^25^ that change the expression of many genes in a coordinated fashion ^26^, and (b) cancer cells attract different immune and stromal cells dilute gene expression measurements with their own regulatory programmes ^27^.

Here, we aim to bridge this gap by investigating how well GRN inference methods perform in the context of cancer, and particularly how much they are influenced by specific confounding factors outside of TF-TG relationships such as aneuploidies and sample impurities.

## 2. Network inference methods have been extensively evaluated on synthetic data sets

### 2.1 Network inference methods

For steady-state networks, the basic idea is that the same underlying regulatory structure (the network to be inferred) will be sampled at different states by measuring gene expression of e.g. multiple cancer patients. Genes that are up- or downregulated in a subset of samples compared to the rest will change their expression in accordance to the underlying regulatory network, which can in turn be inferred by looking at this correlation structure: If two genes are correlated across many samples, they are likely to either regulate each other or be regulated by a common third gene (albeit not always directly).

Classic methods that infer networks from multiple samples of unperturbed gene expression can roughly be divided in correlation-based and information-theoretic models. These and more methods have been reviewed in detail ^8,10,28,29^ and hence we only provide a brief overview. Information-theoretic approaches started out with relevance networks ^12^, in which the pairwise mutual information (MI) is computed between all pairs of genes. Subsequently, all gene pairs above a certain threshold (that can be estimated from the data itself) are kept. ARACNe (algorithm for the reconstruction of accurate cellular networks) ^1,13^ added an additional filtering step where the authors eliminate the weakest link in all gene triplets (using the data processing inequality) unless they are protected by a transcription factor link. The recent ARACNe-AP ^30^ (for adaptive partitioning) implementation adds further performance optimizations. By contrast, the PCIT ^31^ algorithm only removes edges in triplets if two genes are conditionally independent given the third. Other approaches were taken by CLR ^14^ (context likelihood of relatedness; using the z-score of the MI distribution) or C3NET ^15^ (conservative causal core networks; keeping only the strongest MI edge for each gene) and its extension BC3NET ^32^ (bagging of C3NET results). Yet another approach is taken by MRNET ^33^, which concurrently maximizes the relevance (MI) while minimizing redundancy (MRMR is a feature selection technique in supervised learning). For practical purposes, it should be noted that MI-based methods are nonparametric, i.e., these methods perform on the ranks of gene expression values rather than the gene expression values themselves. Many of these methods are implemented in the minet R package ^34^.

Correlation- and regression-based models are another class of gene regulatory network inference methods. In their simplest form, these methods perform a regression or correlation test between two variables. As there are many gene interactions that need to be tested, feature selection is a common feature of these techniques. For instance, Least Angle Regression (LARS) ^35,36^ starts with the best correlated predictor and then iteratively adds other predictors based on their correlation with the residual. TIGRESS (Trustful Inference of Gene Regulation with Stability Selection) ^17^ adds the concept of stability selection to LARS. Instead of adding and removing individual predictors, the GeneNet package ^37^ estimates all predictors simultaneously by inverting the gene expression matrix. Another approach is to combine regression models with decision trees, finding sets of genes that best explain the expression of a target ^38–40^. GENIE3 (Gene Network Inference with Ensemble of trees) ^16^ integrates information of many such trees in order to make regulatory predictions. NIMEFI (Network Inference using Multiple Ensemble Feature Importance algorithms) ^18^ goes one step further and integrates the results of both TIGRESS and GENIE3 into a combined prediction method.

### 2.2 Finding a reference set for method evaluation

A challenge in evaluating network inference methods is that in order to score the performance of different methods, we need to compare the edges they infer to edges we know are correct vs. edges we know not to be correct. However, we often do not know the ground truth for real gene regulatory networks. While many interactions may be known, it is likely that only a small fraction of the relevant interactions has been discovered. An alternative approach is to simulate a GRN according to a known network structure and a set of rules about how the different nodes influence each other. Examples of such simulators are SynTren ^41^ and GeneNetWeaver ^42^. The advantage of such a synthetic network is that the ground truth is known, but it may not exhibit all the properties of a real GRN. Method evaluation was often focused on synthetic or synthetic-like datasets. When evaluations on real data were done, they usually were small in scale owing to the limited amount of orthogonal data available.

More recently, the amount of available human TF-gene interactions has grown tremendously due to large-scale efforts like the ENCODE project ^43^, but also curation of individual ChIP binding experiments ^44^. These have produced consensus regulons for individual TFs, where binding was observed in a variety of tissues. The latter comprise 100 transcription factors that cover 16,500 target genes in a data set available from the Enrichr platform ^45^. Another option would be the UniBind database with 231 transcription factors ^46^. While these consensus interactions are still not proof of actual regulatory interactions, they provide a sufficient number of orthogonally derived relationships in order to use this set to identify large-scale biases of network inference algorithms with respect to copy number changes or sample impurities.

### 2.3 Previously published method evaluations

In terms of previous method evaluations, the most comprehensive benchmark studies are the Dialogue of Reverse Engineering and Assessment of Methods (DREAM) challenges ^11^. These are community-driven challenges where a panel of organizers designs competitions about, among other things, network inference. DREAM4 consisted of five synthetic networks with 100 genes and 100 samples. Each of the 100 genes could be a regulator of other genes. These networks were called “multifactorial”, as all nodes were perturbed simultaneously in the simulations. The expression matrices hence contained 100 different steady state realizations of the same underlying network. DREAM5 ^11^ provided three kinds of networks: a synthetic network, one derived from *E. coli*, and one derived from *S. cerevisiae*. The gene expression matrices were larger and ranged from 1600-5900 genes and 536-805 samples, respectively. In contrast to DREAM4, all networks defined a subset of genes that could act as regulators (between 195 and 334; cf. Fig 1a-b). In addition, newly published methods often perform their own evaluation ^12–14,16–18^.

**Figure 1:**
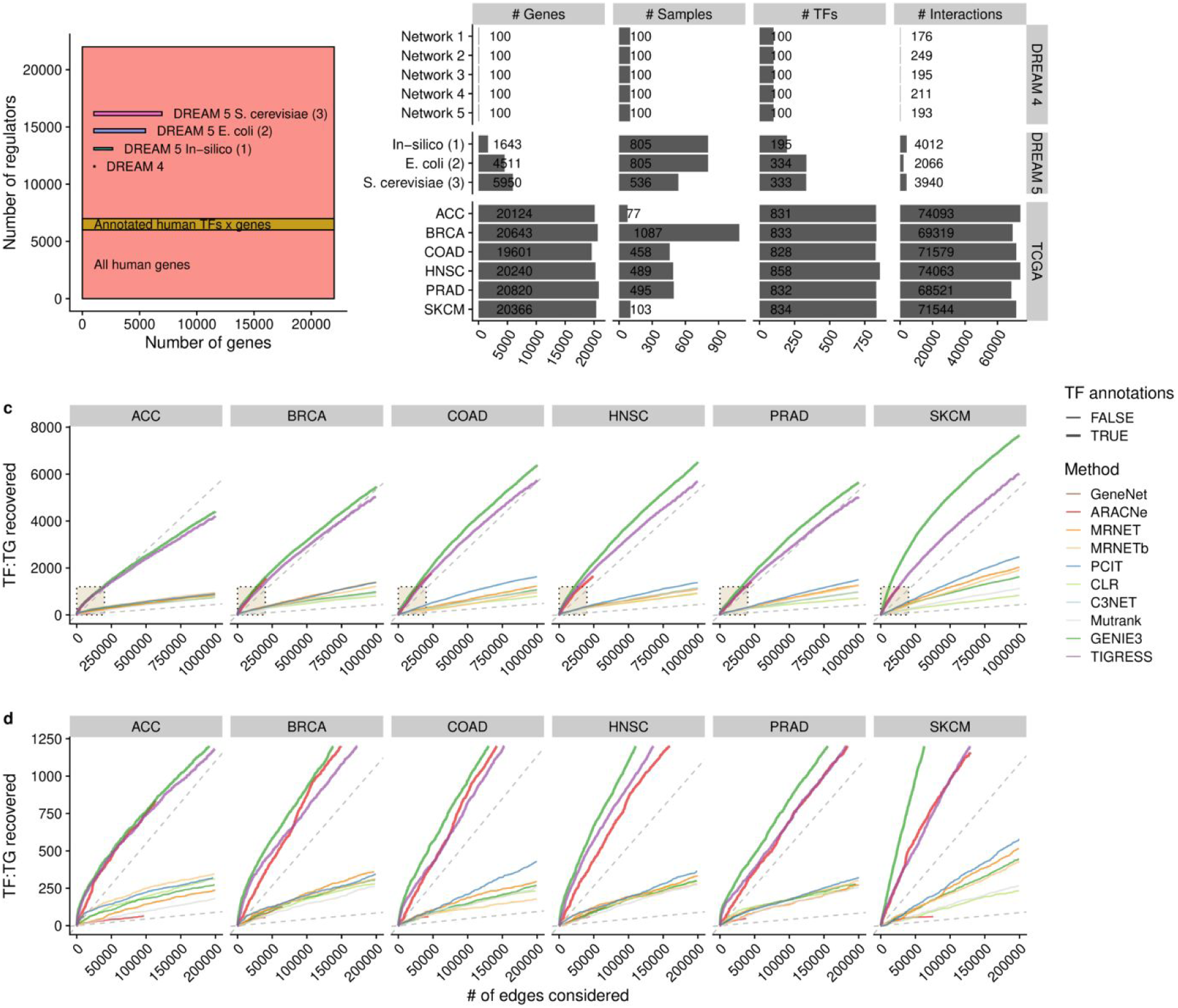
Translating DREAM challenges to a cancer data set. (a) Comparison of potential interactions in the previous challenges vs. all binary interactions in a human genome with and without known regulators. (b) Number of genes, regulators, samples, and known interactions for the different challenges and TCGA cancer cohorts. (c) Number of known TF-TG pairs recovered by network inference algorithms (y axis) for six selected TCGA cohorts with size cutoff of the inferred network (x axis). Dashed lines indicate expected recovery if randomly sampling from all gene interactions (lower line) or only from known regulators (upper line) (d) zoomed view of c for small sizes of the networks.

Yet, research investigating gene regulatory networks is often applied to more complex systems like human cancers ^1,2,19^, and not only to simpler and better-defined synthetic networks or networks from microorganisms. In the context of cancer, a single TF was validated using a set of 26 known targets or comparison between 11 known targets and 11 negative targets ^1^. We are currently not aware of a large-scale study comparing GRN inference methods in the realistic setting of human cancer gene expression ^10^, which may be explained by the fact that it is difficult to obtain a reference network to compare to.

## 3. Previous benchmarks do not accurately capture properties of cancer gene expression

### 3.1 Evaluating inference methods on cancer gene expression data

While the gene expression data sets that were used for evaluation have dramatically increased in size in DREAM 5 compared to DREAM 4, they are still very small in comparison to mammalian organisms (Fig. 1a, b). This starts from the number of genes and regulators present, but is also apparent by the number of confirmed regulatory interactions in the DREAM 5 networks. By contrast, the number of samples available is often not higher than in the much simpler benchmark data set (Fig. 1b). Hence, real cancer gene expression data offers a different kind of challenge for inference methods and may lead to different results compared to the previously published benchmarks. However, as the same methods are commonly employed to infer regulatory programmes in cancer, it is important to gain a better understanding of the opportunities and pitfalls that are specific to this kind of data and may not have been accurately covered in simulation studies or the other DREAM benchmarks.

Here, we are not only interested in the more complex system as defined by the number of genes and regulators, but also in specific biases that cancer gene expression exhibits and that was not covered sufficiently by synthetic or micro-organism networks, like copy number alterations or sample mixtures due to stromal and invading immune cells. As the network simulators covered ^41,42^ do not allow for these kinds of biases, we aim to evaluate GRN inference methods on cancer patient gene expression from The Cancer Genome Atlas (TCGA) ^47^.

We chose six TCGA cohorts with different numbers of samples available: Adrenocortical carcinoma (ACC), Breast invasive carcinoma (BRCA), Colon adenocarcinoma (COAD), Head and neck squamous carcinoma (HNSC), Prostate adenocarcinoma (PRAD), and Skin cutaneous melanoma (SKCM). These range from 77 (ACC) to 1087 (BRCA) samples per tumor type (cf. Fig. 1b). We filter all mapped genes to those with 5 or more reads on average per sample, yielding approximately 20,000 genes for all cohorts. We define potential regulators as genes that are annotated with “Transcription factor activity” in Gene Ontology ^48^ (GO:0003700), leaving approximately 830 regulators per cohort. As a positive set, we used consensus regulons for 101 transcription factors covering 16,500 target genes from ChEA ^44^ and ENCODE ^43^ via the Enrichr platform ^45^. We did not use a negative set, as those are generally not available for transcriptional regulation.

We ran the network inference algorithms ARACNe-AP ^30^, GeneNet ^37^, GENIE3 ^16^, TIGRESS ^17^, and other methods available via the NetBenchmark R package ^49^ on each of our six cohorts using default options. We then evaluated how many known TF-TG pairs were recovered in the top *N* edges (Fig. 1c and 1d), prioritized by the score given to each interaction by each method. We found that GENIE3 has a slight edge over TIGRESS, which again performs slightly better than ARACNe-AP. The difference between these three methods and all others is that they take into account TF annotations, which the other methods do not.

### 3.2 Incorporating prior knowledge is essential for method performance

In terms of network size, the number of potential interactions with knowledge about regulators compared to without is particularly striking: If any gene can also act as a regulator, there are approximately 22,000 genes and 484 million binary interactions. By contrast, using 974 annotated TFs in Gene Ontology ^48^ and only taking them into account as potential regulators, we are left with only 21 million potential interactions to explore (a 22 fold decrease; cf. Fig. 1a). However, note that all the networks that we infer are undirected, hence these numbers should be halved when considering how well a method recovers known binding interactions. Also, we do not allow self-regulation, i.e. edges of a gene with itself.

This decrease in potential interactions seems to drive a superior performance of methods that are able to incorporate TF annotations in the network they infer. No matter the background or the age of a method, looking naively at the number of links recovered from known ChIP binding, there is a substantial increase (Fig. 1c, d). This result should of course be regarded with respect to the number of potential interactions: sampling randomly from all gene-gene interactions will produce a much worse performance than sampling from known TFs (lower and upper dashed grey lines in Fig. 1c, d, respectively). Nevertheless, if we are interested in recovering true regulatory interactions, it stands to reason to use a method that makes use of TF annotations. If not, no method ignoring these annotations performs anywhere close to randomly sampling from TF-TG interactions (upper dashed line in c, d).

It should be noted that the performance of all methods are close to their respective random lines. This is likely explained both by the fact that our positive set likely contains many non-regulatory binding interactions, and the observation that none of the methods in DREAM 5 performed much better than random for the *S. cerevisiae* network ^11^. However, for network sizes up to 200,000 nodes GENIE3, TIGRESS, and ARACNe-AP perform better than random sampling of TF-TG interactions (cf. Fig. 1d).

### 3.3 Copy number changes and sample impurities are confounding gene expression measurements

As gene regulatory networks are often inferred from gene expression, it is important to consider factors influencing gene expression outside of transcription factor-target gene relationships. This is why care needs to be taken when merging together multiple data sets from e.g. microarrays and RNA-seq, or different processing pipelines that can lead to technical batch effects. These batch effects have been abundantly discussed in literature before (reviewed in ^50^), and there are many approaches to correct for them ^51–53^.

However, cancer cells also harbor biological variability influencing gene expression and hence correlation that has so far not been discussed in depth. For instance, cancer cells often harbor copy number changes ranging from small segments (focal CNAs) up to the level of whole chromosomes (aneuploidies) ^54^. Gene expression has been shown to follow these copy number changes ^26,55,56^, whereas protein expression is often compensated for ^57^.

Another factor that influences cancer gene expression in particular is that samples obtained from patients will not only consist of a homogeneous population of cancer cells. Instead, samples will also contain stromal cells that have been co-opted in tumorigenesis, as well as immune cells ^27^ driving inflammation and/or contributing to active clearing of tumor cells. Multiple methods have been developed to estimate cell fractions ^58–61^, including some that aim to reconstruct the cancer-specific transcriptome from a cell mixture ^62,63^. Another level of complexity is that cancer cells themselves often consist of multiple clones and lineages that may exhibit heterogeneous traits not visible in a bulk transcriptomics measurement. While these issues can be overcome with recent single-cell sequencing technologies, it will still take years until these data sets reach a level of comprehensiveness comparable to the TCGA.

Here, we focus both on focal copy number changes and aneuploidies, as well as tumor purity defined as the fraction of cells in a sample estimated to be cancer cells. As focal copy number changes, we take recurrently altered regions (RACS) from the Genomics of Drug Sensitivity in Cancer (GDSC) project ^64^ as processed by ADMIRE ^65^. For different cancer types in the TCGA, we observe a different number as well as different sizes for these regions (Fig. 2a). Glioblastoma multiforme (GBM) and Ovarian serous cystadenocarcinoma (OV) show the highest number of altered regions, with Breast invasive carcinoma (BRCA) and Lung squamous carcinoma (LUSC) showing the highest fraction of their genomes altered due to these local recurrent events. We calculate aneuploidy scores as the average absolute deviation from euploid over whole chromosomes according to copy number segments downloaded via TCGAbiolinks ^66^ (Fig. 2b), and use consensus purity estimates for different samples from the xCell publication ^59^.

**Figure 2:**
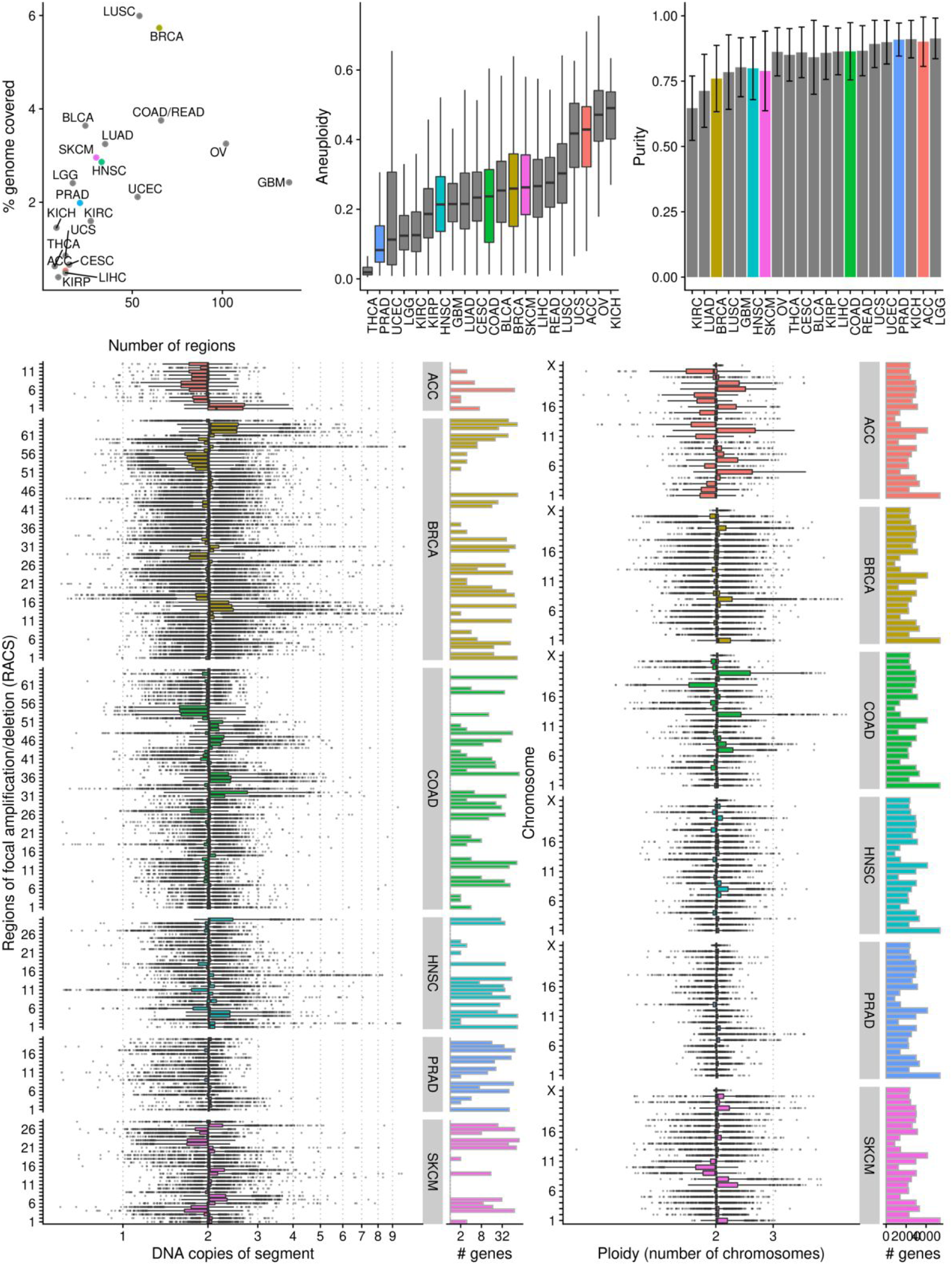
Abundance of focal amplifications and aneuploidies in the TCGA. (a) Number of recurrently altered focal segments (x axis) vs. the fraction of the genome that they cover (y axis) for different TCGA cohorts. (b) Distribution of aneuploidy scores for different TCGA cohorts with chosen cohorts highlighted in color. (c) Average sample purity by cohort, error bars are standard deviation (d) Distribution of segment (left) and chromosome (right) copy numbers across samples of the six chosen cohorts.

The cohorts we focus on in subsequent analyses show a heterogeneous level of focal copy number changes and aneuploidies, as well as for sample purity (Fig. 2a-c). Looking at individual samples, we can observe the variability in focal amplifications from an almost euploid cohort (PRAD) up to a very high level (BRCA; cf. Fig. 2d). Similarly, PRAD also shows a low and ACC a high level of aneuploidy (Fig. 2e).

In terms of gene regulatory networks, it is unclear how these factors confounding gene expression influence the inferred edges for different methods. This is why in this study we evaluate the number of edges each of our methods infers that fall within (1) a CNA vs. outside and (2) genes whose expression strongly correlates with sample purity vs. those that do not. As a control, we check for the same enrichment in known transcription factor binding sites.

## 4. Network inference methods are biased towards copy number aberrations and sample purity

### 4.1 Focal amplifications have strong local but weak genome-wide effect

In order to test for the effect that focal amplifications have on the interactions inferred by different network inference methods, we used our selected methods and cohorts to investigate how many of the inferred edges can likely be explained by the focal amplifications and aneuploidy scores that we previously obtained (Fig. 2). Briefly, we assume that real TF-TG relationships (obtained from ChIP binding information) are equally likely to occur within focal CNAs or aneuploidies as they are between two genes anywhere on the genome. We then compare the edges obtained by the different inference methods to the number of edges theoretically possible within CNAs, and see if this fraction is different to the total number of edges inferred from the total number of possible edges. As a control, we do the same for known transcription factor binding associations to confirm our assumption of equally likely TF-TG relationships within and outside of CNAs.

Because all of the methods we tested provided a score for each inferred edge, we could test different network sizes by setting different cutoffs on the edge scores. We then went on to show the number of expected false positive edges with different network sizes for our six highlighted cohorts (Fig. 3). As our positive set that we test with is likely incomplete, we can only estimate the fraction of these false positive links, and not if any individual link is indeed a false positive.

**Figure 3:**
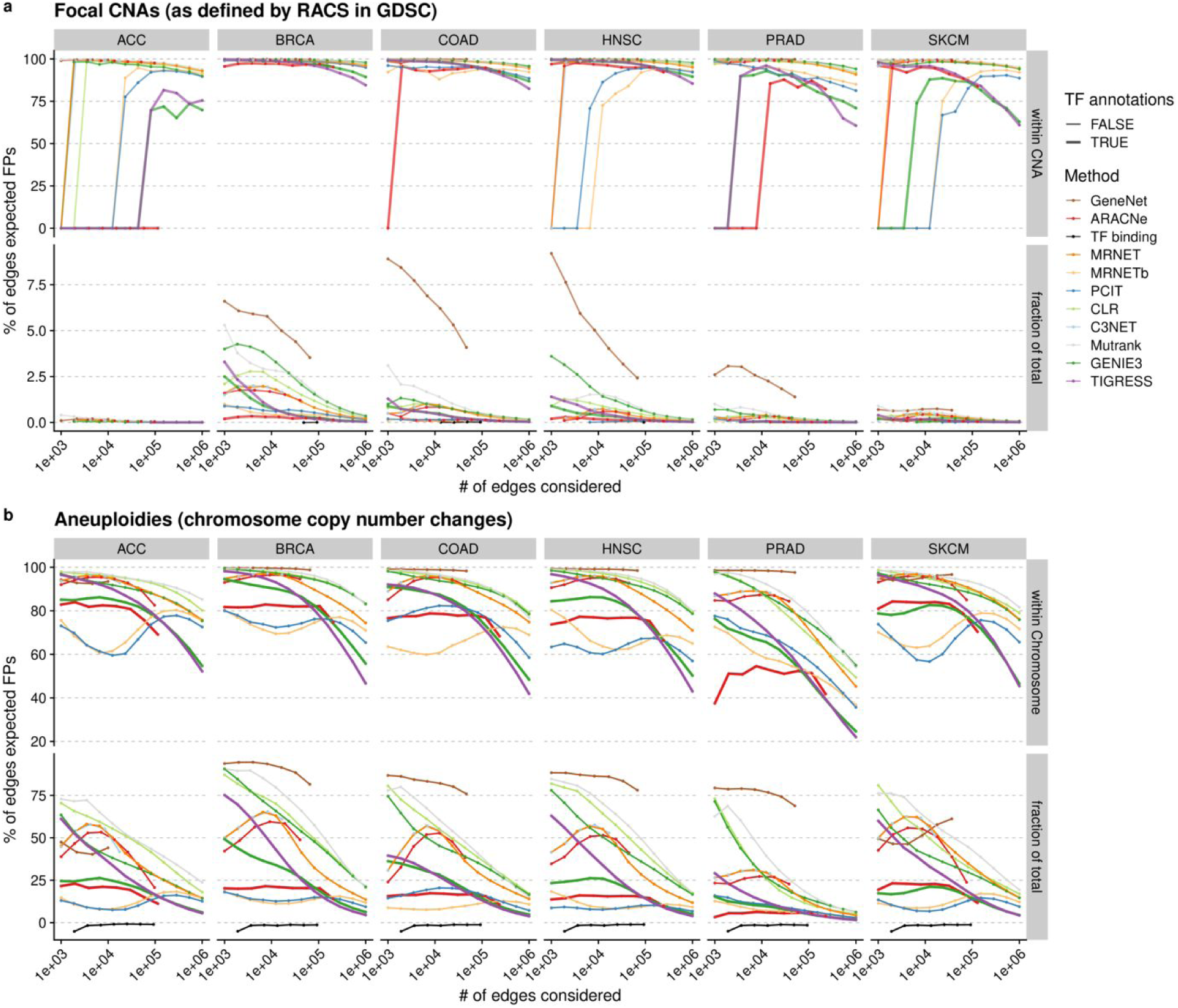
Effect of (a) focal amplifications and (b) aneuploidies on inferred network edges. False positive rate (y axis) shown for different network size (x axis) either within the CNA (top row) or in comparison to the total number of edges between all genes (bottom row). Dots along lines are shown where this was quantified.

First, we show the effect of focal amplification on the inferred edges (Fig. 3a). On the top row, we show the sum of observed links within each CNA over the number of possible links within those CNAs. We find that starting from very small networks (1000 edges), almost all of the within-CNA edges inferred by different methods in most cohorts are likely false positives (FPs), as we observe many more edges than we could expect given our null model (that edges within a CNA and outside are equally likely). These false positives, however, have got a relatively minor impact on all the edges inferred across the genome, as the number of genes in the identified recurrent focal amplifications is low (cf. top vs. bottom row in Fig 3a).

GeneNet is the method that shows the strongest enrichment of edges in CNAs, with up to 10% of the total number of edges in a small network (1000 edges). Other methods stay under 5% of genome-wide false positive edges. As the size cutoff gets less stringent (100,000 to 1 million edges), the genome-wide FPR for most of the methods and cohorts drops under 2%. It should, however, be noted that the total number of possible within-CNA edges is low for all cohorts and incorporating all of them in a network will still result in a low genome-wide FPR (cf. Fig 2d). Hence, a more relaxed definition of recurrent focal CNAs (compared to the one defined by ADMIRE) would likely also yield a higher rate of genome-wide FP edges.

### 4.2 Aneuploidies have weaker local but strong genome-wide effect

In contrast to the focal amplifications, aneuploidies show a smaller within-segment FPR (Fig. 3, top rows). The effect of the genome-wide FPR, however, is much bigger for aneuploidies than for focal amplifications (Fig. 3, bottom rows). This makes sense intuitively, as the number of genes changed with each aneuploidy is much larger than the number of genes changed with a focal amplification (cf. Fig. 2c-d). Hence, a smaller fraction of incorrectly identified edges within each chromosome already has a large effect on the genome-wide false positive rate. We can observe this in the FP curves between focal amplifications (Fig. 3a) and aneuploidies (Fig. 3b) that reach a much higher level of genome-wide FPR for aneuploidies (up to 85% of the total number of edges inferred, compared to under 10% for focal amplifications).

These results suggest that aneuploidies are likely a major source of bias in the total number of inferred edges for most methods, especially for smaller network sizes. Therefore, caution should be taken when applying these methods to biological samples that may harbor large-scale copy number changes. For the methods that performed well in recovering TF-TG interactions (cf. Fig. 1c-d), we see that TIGRESS is most influenced by aneuploidies, followed by GENIE3 (although the effect is reversed in melanoma). Both methods show a larger FP enrichment with smaller network size, suggesting that they are prone to assigning high scores to genes co-regulated by aneuploidies instead of TF-TG interactions. ARACNe is remarkably stable in the fraction with varying network size, always showing approximately 10-20% of FPs due to aneuploidy. All methods using TF annotations behave equally in the range of 100,000 to 1 million edges. Actual TF binding interactions from ChEA (black line in Fig. 3) are equally likely to be inside and outside of CNAs.

### 4.3 Networks are biased due to sample composition

Apart from the copy number aberrations, we also investigated the number of false positive edges with sample impurities. The rationale is the same as above: we assume that genes whose expression level is changing with tumor purity are not more likely to be transcription factor and target gene compared to the genes whose expression does not follow that trend. A difference to the analysis before, however, is that there is no clear set of genes that are influenced by sample purity vs. genes that are not. In order to address this, we selected either the top 1,000 or top 5,000 genes that correlated the most with tumor purity in each cancer type we highlight, and then performed the same enrichment analysis as we did with the focal copy number changes and aneuploidies.

For the methods that performed well in recovering TF-TG interactions, we see that GENIE3 is more susceptible to wrongly inferring links due to samples mixtures than TIGRESS, with more FPs in smaller networks (10-40% vs. 10-20%, respectively). ARACNe shows a stable FPR of 10-15% irrespective of network size. From 100,000 to 1 million links, the methods largely equalize. Again, we do not observe an enrichment of TF binding with purity-correlated genes (black line in Fig. 4). As with the results we find for the CNAs, there is a trade-off between the within-chromosome FPR vs. the genome-wide FPR depending on the cutoff for selecting the purity-correlated genes (albeit less pronounced).

**Figure 4:**
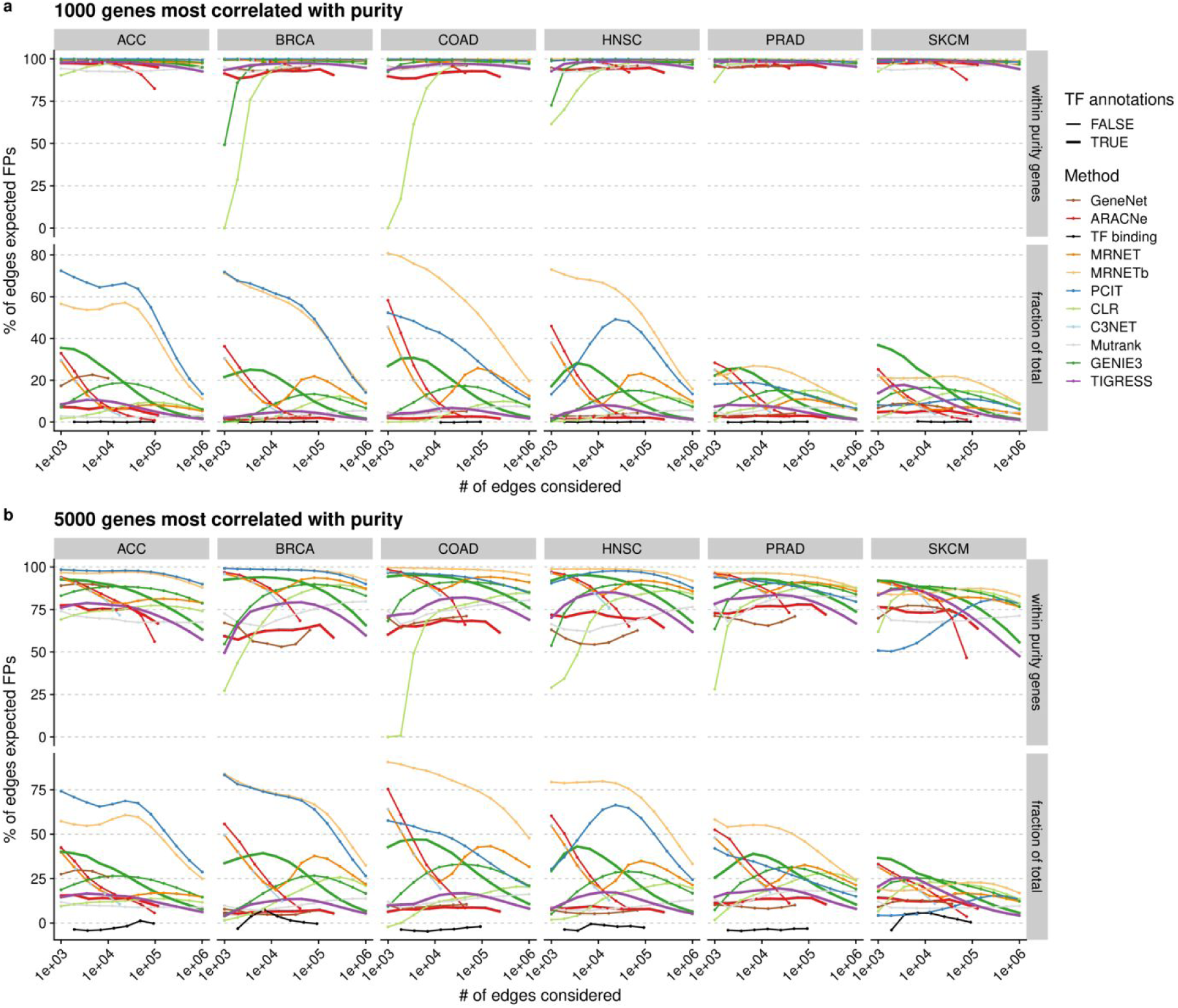
Effect of sample purity on inferred network edges. For (a) 1000 or (b) 5000 genes most correlated with sample purity, expected false positive rate (y axis) is shown for different network sizes (x axis) in the chosen cohorts.

## 5 Conclusion

When inferring gene networks in the context of cancer, it is important to not only keep in mind the potential technical variability between batches that may induce false positive correlations and hence edges when using network inference methods, but also the biologically intrinsic confounding factors of gene expression, like the one induced by DNA copy number changes ^26,55^ or a mixture of different proportions of different cell types ^60,61^.

We have shown that for recovering an accurate network of TF-TG interactions in cancer, methods that incorporate TF annotations should be preferred to those that are not. However, even these methods are largely susceptible to inferring false positive links due to confounding factors. Combined, aneuploidies and sample impurities can be expected to contribute approximately 20-30% of false positive edges for networks with 100,000 to 1 million edges, and a larger fraction for smaller networks.

If we are interested in accurately inferring true regulatory interactions, there is a need to correct for these biases. Previous methods have been proposed to correct for confounding factors in gene networks in general (like Principal Component Analysis ^67^ and multivariate linear models ^68^), but they have not investigated how well their methods adjust for biological influences like the ones we discussed here. In addition, future method development in this area could address these biases more directly by also modeling aneuploidies and sample impurities explicitly.

## Online methods

### Gene expression and copy number data from the TCGA

We have downloaded the raw read counts for gene expression as well as the inferred continuous regions of the same DNA copy number (copy number segments) from the harmonized TCGA data obtained through the R package TCGAbiolinks ^66^. We chose the cohorts (ACC, BRCA, COAD, HNSC, PRAD, and SKCM) because they represented a wide range of sample sizes, ploidy, and purity values (cf. Fig 2).

We further filtered the samples set to only contain primary tumors (TCGA sample type of “01A”). We filtered the genes to only contain genes on human chromosomes 1 to 21 (excluding X, Y, and MT) and to have more than 20% of the samples with 10 or more reads. We then used the DESeq2 R package ^69^ to estimate library size factors and get variance stabilized gene expression values.

Finally, we mapped Ensembl IDs to HGNC gene symbols using Ensembl 96 and removed all genes that did not have a valid gene symbol or were duplicated.

### Focal and chromosome copy numbers

For regions with recurrent copy number alterations for both our cohorts, we downloaded Table S2D from https://www.cancerrxgene.org/gdsc1000/GDSC1000_WebResources/^64^.

To get copy numbers of either these segments or whole chromosomes, we calculated the average copy number along the respective regions for each sample in our cohorts.

### Sample purity and purity-correlated genes

We obtained consensus purity estimate from xCell ^59^ using their “estimate” field in their Additional File 6.

We then, for the primary samples of each of our cohorts, calculated differential expression along this estimate (using a likelihood ratio test over the intercept using the DESeq2 package), and selected the top *N* genes by lowest p-value.

### Transcription factor annotations and binding data

For genes that may act as transcription factors, we downloaded all HGNC symbols associated with GO:0003700 (DNA-binding transcription factor activity) from Ensembl 96 ^70^.

For our positive set of real transcriptional regulation, we downloaded the “ENCODE_and_ChEA_Consensus_TFs_from_ChIP-X” HGNC symbols from the Enrichr platform ^45^, encompassing 16,500 different target genes of 100 transcription factors.

### Network inference

For inferring our networks, we used the following methods: ARACNe-AP ^30^, which is a Java implementation that extends the original ARACNe method; the GeneNet R package ^37^; the GENIE3 R package ^16^, the TIGRESS R package ^17^, and other methods available in the NetBenchmark R package ^49^.

We infer one network per cohort per method, and look for enrichment of edges that are likely due to copy number changes or sample mixtures. As not all of these methods provide a significance measure, we instead look at the order of edges inferred, from the highest score to the lowest.

### Quantifying possible TF-TG interactions

To quantify enrichment of real (obtained from ChIP binding experiments) TF-TG interactions within the top *N* genes of a given network, we first need to enumerate the possible number of edges given how many genes and transcription factors we have, and whether a GRN inference method knows the difference between regulators and targets.

In particular, if there are no known regulators we consider the number of possible edges to be:

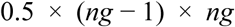

And if they are known instead:

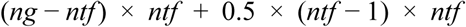

Where *ng* is the total number of genes and *ntf* is the number of potential regulators. Note that if every gene can be a regulator, the lower formula simplifies into the upper.

### Enrichment of edges within gene sets

We quantify bias by copy number changes or purity by assuming that the genes in a set (focal regions, chromosomes, or purity-associated genes) are equally likely to form links within the respective set as they are with genes outside the set. When then look for enrichment of edges within a given region over the total number of edges.

In particular, we first compute the odds ratio of a method obtaining links in a segment vs. the overall number of edges:

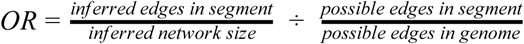

Then, the local false positive rate (FPR) is defined as:

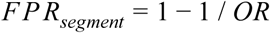

While the genome-wide FPR is the number of expected FP links divided by the size of the inferred network:

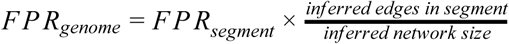

## Code availability

The analysis code of this manuscript, including code to generate all figures is available at https://github.com/mschubert/GRN-aneup-purity, licensed under GNU GPL version 3 or later.

